# Controlling for effects of confounding variables on machine learning predictions

**DOI:** 10.1101/2020.08.17.255034

**Authors:** Richard Dinga, Lianne Schmaal, Brenda W.J.H. Penninx, Dick J. Veltman, Andre F. Marquand

## Abstract

Machine learning predictive models are being used in neuroimaging to predict information about the task or stimuli or to identify potentially clinically useful biomarkers. However, the predictions can be driven by confounding variables unrelated to the signal of interest, such as scanner effect or head motion, limiting the clinical usefulness and interpretation of machine learning models. The most common method to control for confounding effects is regressing out the confounding variables separately from each input variable before machine learning modeling. However, we show that this method is insufficient because machine learning models can learn information from the data that cannot be regressed out. Instead of regressing out confounding effects from each input variable, we propose controlling for confounds post-hoc on the level of machine learning predictions. This allows partitioning of the predictive performance into the performance that can be explained by confounds and performance independent of confounds. This approach is flexible and allows for parametric and non-parametric confound adjustment. We show in real and simulated data that this method correctly controls for confounding effects even when traditional input variable adjustment produces false-positive findings.

## INTRODUCTION

Machine learning predictive models are now commonly used in clinical neuroimaging research with a promise to be useful for disease diagnosis, predicting prognosis or treatment response (Wolfers et al. 2015). They are also being used in non-clinical settings to detect possible relationships between biology and personal characteristics such as cognitive capabilities, or identify neural correlates of stimuli or a task (Naselaris et al. 2011). For the correct interpretation of the results and translation of machine learning models into clinical practice, it is important to verify that the machine learning predictions are not driven by the effects of confounding variables. For example, in a cognitive experiment, accurate predictions of a stimulus identity can be caused by head motion or increased effort due to task difficulty, instead of a neural signal of interest. In a clinical setting, gender, scan-site, motion, or age can cause seemingly accurate machine learning prediction, capturing no other useful information about the disease.

The most common way to control for confounds in neuroimaging is to adjust input variables (e.g., voxels) for confounds using linear regression before they are used as input to a machine learning analysis (Snoek et al. 2019). In the case of categorical confounds, this is equivalent to centering each category by its mean, thus the average value of each group with respect to the confounding variable will be the same. In the case of continuous confounds, the effect on input variables is usually estimated using an ordinary least squares (OLS) regression. The input variables are adjusted by subtracting the estimated effect (i.e., taking the residuals of the confound regression model). This method is, however, problematic for confound adjustment for machine learning models. Since machine learning models are often non-linear, multi-variable, and not fitted using OLS, they can extract information about confounds that OLS regression does not remove. Thus, even after confound adjustment of input variables, the machine learning predictions might still be driven by confounds.

We propose an alternative approach to control for confounds at the level of machine learning predictions instead of the level of input variables. This avoids the problems of input adjustment because we only need to estimate the effect of confounds on the outcome, so that it produces valid results even for complex machine learning models. This method has an intuitive interpretation: estimating the proportion of variance in the outcome can be explained by model predictions that are not already explained by confounding variables.

The remainder of this paper is organized as follows: First, we illustrate multiple circumstances where adjustment of input variables does not sufficiently control for confounding effects. Second, we describe our proposed method, including important caveats relating to the adjustment in regression and classification settings, non-linear and non-parametric adjustment, and cross-validation and permutation schemes. Third, we will show using simulated and real data that the proposed method adequately controls for confounds, even in situations where traditional input adjustment fails. Last, we will discuss how our work relates to other methods, and to which extent published results can be affected by insufficiently adjusted confounds.

## PROBLEMS OF INPUT ADJUSTMENT

Confound adjustment of input variables is the most widely used method for controlling for confounds in neuroimaging machine learning studies. It is based on a simple idea that if we remove the effects of confounds from the input variables, the results of the subsequent analysis will not be affected by confounds. The effect of confounds is usually estimated using an ordinary least squares (OLS) linear regression for each input variable separately:

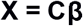

where X is a vector of the to be adjusted input variables (e.g., voxels or regions of interest), C is a matrix of confounds, and B is a vector of regression coefficients estimated using OLS. The variable is then adjusted by subtracting the estimated effect of confounds

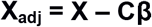

However, this method does not guarantee that the subsequent machine learning analysis will not be affected by confounds. This is because machine learning models can capture information in the data that cannot be captured and removed using OLS. Therefore, even after adjustment, machine learning models can make predictions based on the effects of confounding variables.

There are several sources of confounding information that the OLS adjustment method cannot remove. These are illustrated schematically in Figures 1 and 2 in the context of a machine learning classification and regression, respectively. These plots show scenarios where only confounding variables are added to the data (i.e. no signal) which are then regressed from the data using OLS. Nevertheless, a classifier (Figure 1) or regression model (Figure 2) can still detect signal. First, usually, only linear effects are removed, but nonlinear effects will still be present in the data. This can be mitigated by fitting a more complex model using, for example, regressions with polynomial or basis spline expansion. However, even with a complicated model, it is not guaranteed that the model fits the data well. In traditional GLM analysis, this could be easily checked using, for example, diagnostic residual plots. However, such a manual check is not feasible for the large number of variables commonly included in neuroimaging machine learning studies.

**Figure 1:**
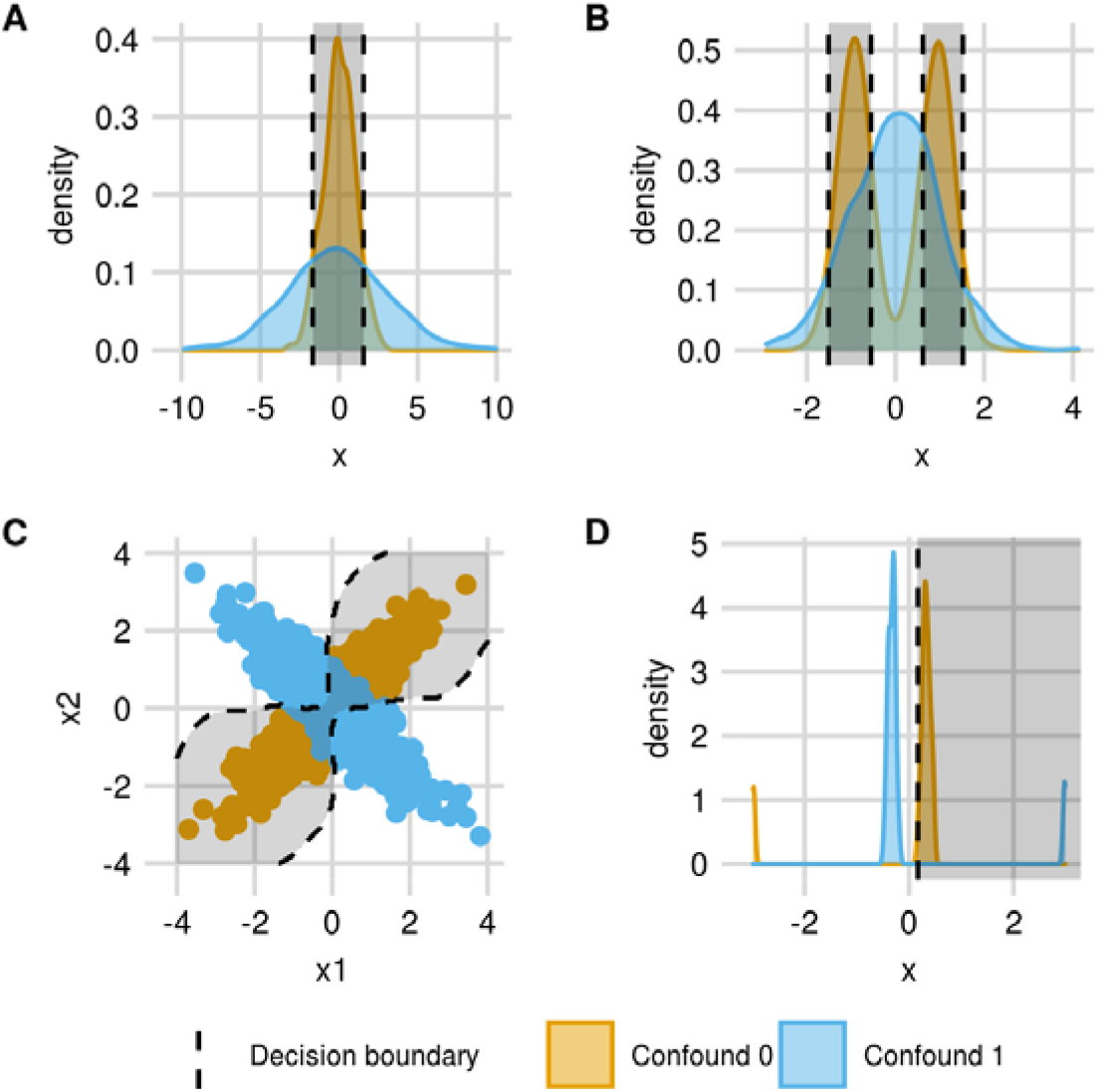
Examples of classification situations where confounding variables are still discriminable using machine learning methods even after regressing out confounds from input variables. **A**: The confound does not affect the mean, but it affects the variance, which can be learned by a linear or nonlinear machine learning model. **B**: Two groups have the same mean, same variance, but different shape of the distribution, which can also be picked up by a machine learning model. **C**: The individual variables x1 and x2 have the same marginal mean, variance, and shape, however, their joint distribution is different between two groups, which can be detected by a nonlinear machine learning model. **D**: Two groups have the same mean, however, due to outliers, a robust machine learning model can still discriminate between the two groups.

**Figure 2:**
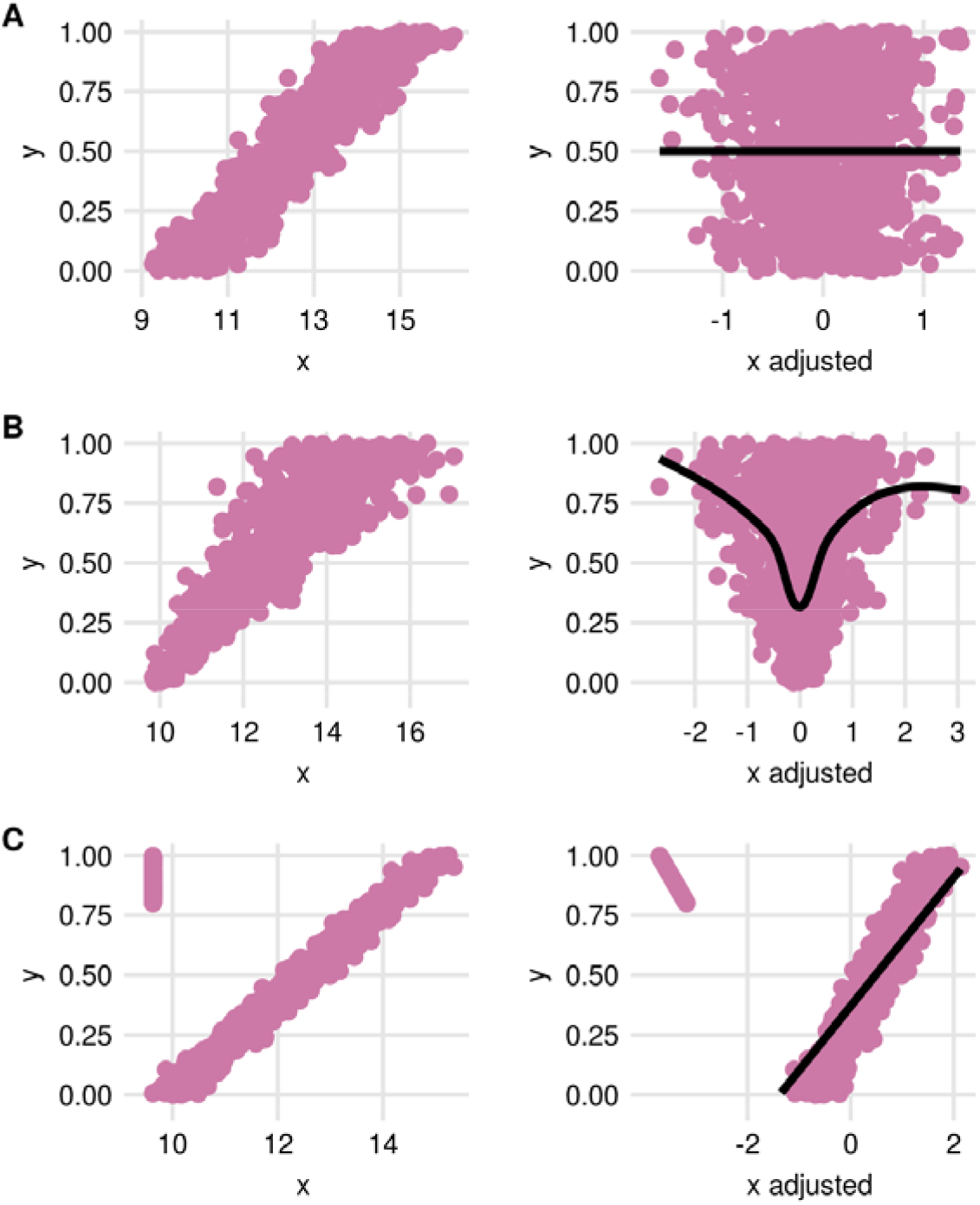
Examples of situations when adjusting input variables for continuous confounding variables does and does not work. Left: original variable before adjustment. Right: variables after confound adjustment. **A:** A situation where confound adjustment works. A linear effect of y on x is removed, and there is no more information about y left in the data. **B:** there is heteroskedastic noise in the input variables. Although the linear effect is removed, there is still confounding information left in the data that can be picked up by a nonlinear machine learning model. **C:** A situation where confound adjustment does not work due to a different loss function. A linear effect is removed from the data as estimated by ordinary least squares regression, which is affected by outliers. A robust machine learning model (in this case support vector machine) can learn a relationship between confound and outcome variable even after adjustment.

Second, the confounds can affect the scale or shape of the data distribution. For example, in a multi-site analysis, the data variance might be higher in data from one scan-site than another. As was described by Görgen and colleagues (2017), differences in variance can be learned by non-linear but also linear machine learning models. Therefore, even after centering by site, a machine learning model can learn that subjects from one site are more likely to have extreme values of input variables than subjects from the other site (Fig 1A, Fig 2B). This can be mitigated by additionally adjusting the scale of the residuals. The simplest way is to divide residuals in each scan site by their standard deviation or model the residuals' standard deviation as a random effect. Such a modeling approach is performed by ComBat procedure for adjustment of batch effects of microarray data (Johnson et al. 2007) and scan-site effects of MRI data (Fortin et al. 2017). However, this will not help if the confounds affect not only the scale of the distribution but also its shape, such as skewness or kurtosis (Fig 1b). Third, confounds might have a multivariate effect or they may affect the interaction between input variables. Since each variable is adjusted separately, it is impossible to remove multivariate effects, although they can be easily captured using nonlinear machine learning models (Fig 1C). Since OLS regression is fitted to minimize mean squared error, machine learning models that do not minimize mean squared error might still be able to capture confounding information from the data (Fig 1D, Fig 2C). The most prominent example is SVM, which minimizes the hinge loss instead of mean squared error. The hinge loss is less affected by outliers that mean squared error, thus if outliers are present in the data, OLS will be heavily influenced by these outliers whereas robust machine learning models will still be able to capture information about confounds from the adjusted data.

## PROPOSED METHOD TO CONTROL FOR EFFECTS OF CONFOUNDING VARIABLES

We propose that the machine learning predictions themselves should be controlled for confounds instead of individual input variables. We treat machine learning predictions as we would any other potential biomarker and apply traditional regression techniques for confound adjustment (multiple linear and logistic regression) (Pourhoseingholi et al. 2012). This approach aims to estimate, after the machine learning model is fitted, what proportion of variance is explained by machine learning predictions that cannot be explained by confounds.

Using model predictions as an input to an additional regression model to evaluate its performance is not a new idea; it goes back at least to Smith and Rose (1995). The proposed approach is closely related to a method known as pre-validation (Tibshirani and Efron 2002; Hoffling and Tibshirani 2008) used in microarray studies to test if a model based on microarray data adds anything to clinical data. In this section, we will first focus on the most general problem of confound adjustment for machine learning regression and machine learning classification in an independent test set. Next, we will describe the usage of this approach when the machine learning model is evaluated using cross-validation and permutation testing. Last, we will describe non-linear and non-parametric methods for confound adjustment and choice of subjects for creating the adjustment model.

## CONFOUND ADJUSTMENT IN AN INDEPENDENT TEST SET

In a regression setting, there are multiple equivalent ways to estimate the proportion of variance of the outcome explained by machine learning predictions that cannot be explained by the effect of confounds. One is to estimate the partial correlation between model predictions and outcome controlling for the effect of confounding variables.

Given n-dimensional vectors of outcome values **y**, machine learning predictions **p** and a matrix of confounding variables **C**, the partial correlation is equivalent to the correlation coefficient between residuals of **p** and residuals of **y** after regressing out confounds from both **p** and **y** (Whittaker 1990; Kim 2005), that is:

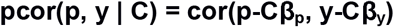

where **β_p_** and **β_y_** are regression coefficients estimated using OLS from a regression equation

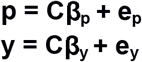

where **e_p_** and **e_y_** are vectors of error values.

The statistical significance of the partial correlation can be obtained parametrically using a Student‘s t distribution (Sheskin 2000; Kim 2005) where the t-statistic is calculated as

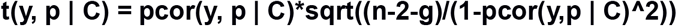

where n is the sample size, and g is the total number of confounding variables. The statistical significance of the partial correlation is equivalent to fitting a linear regression model with machine learning predictions **p** and confounds **C** as covariates and testing if, in this model, **p** is statistically significant, or if adding **p** into the model predicting **y** based on confounds **C** will significantly improve R^2^ using an F test.

For the interpretation, we can decompose the proportion of explained variance to the variance explained only by confounds (**ΔR^2^_c_**), variance explained only by machine learning (ML) predictions (Δ**R^2^_p_**), and variance explained by both confounds and ML predictions (**R^2^_p∩c_**).

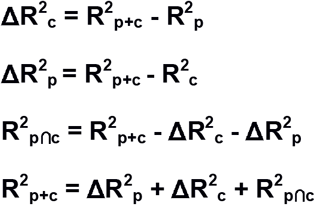

Where **R^2^_p+c_**, **R^2^_p_**, and **R^2^_c_** are **R^2^** of models containing ML predictions and confounds, ML predictions, and confounds, respectively.

For categorical outcomes (classification), the logic of the procedure is the same, but instead of performing adjustment using linear regression, we use logistic regression. Our goal is the same as for regression. We want to know what information about the outcome we can explain using model predictions that is not already explained by confounding variables. This can be similarly expressed as the added value of **p** in the model that includes confounds. The optimal way to do this in a logistic model is to evaluate likelihood ratio (LR) or difference in log-likelihoods of two models: 1) predicting outcome using confounding variables

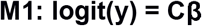

and 2) predicting the outcome using confounding variables and ML predictions.

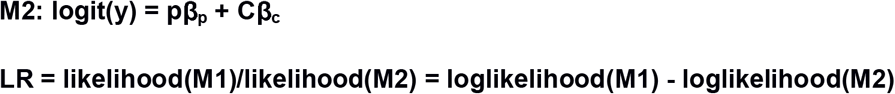

It is important to point out that – similar to the regression setting – this procedure ignores potential miscalibration of predictions, such as systematic overconfidence or underconfidence of estimated probabilities. Thus, we are testing if model predictions contain any information about the outcome that is not already contained in the confounds, but not if the machine learning model's absolute prediction error is better than that of the model using only confounds as predictors. Statistical significance of the partial correlation and likelihood ratio test statistics can be computed parametrically or non-parametrically using a permutation test.

Instead of variance explained, which is not a meaningful measure of model fit for a categorical outcome, we can use a fraction of deviance explained **D^2^**, also known as **R^2^_kl_** due to its connection to Kullback-Leibler divergence (Cameron and Windmeijer 1995). This is equivalent to a fraction of variance explained in linear regression, and in logistic regression, it can be interpreted as a proportion uncertainty reduced due to the inclusion of variables to a model (Cameron and Windmeijer 1995). Another benefit of this measure is that it is closely related to the likelihood ratio test that we use to test the added benefit of ML predictions.

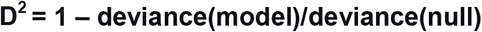

where deviance is **2*(loglikelihood(saturated_model) – loglikelihood(model))**, the *s*aturated model represents a model with one parameter per observation, and null is an intercept only model. The saturated model is a model that has a perfect fit to the data and thus achieves the maximum possible log-likelihood for the given dataset (McCullagh and Nelder 1989).

**D^2^** can be decomposed into a **ΔD^2^_c_ ΔD^2^_c_** and **D^2^_p∩c_** in the same way as **R^2^** as explained before.

## CONFOUND ADJUSTMENT IN CROSS-VALIDATION AND PERMUTATION TEST

The parametric computation of the statistical significance is only valid when the machine learning model is evaluated in an independent test set. However, if a machine learning model is evaluated in cross-validation, traditional parametric tests will produce overly optimistic results. This is because individual errors between cross-validation folds are not independent of each other since when a subject is in a training set, it will affect the errors of the subjects in the test set. Thus, a parametric null-distribution assuming independence between samples will be too narrow and therefore producing overly optimistic p-values. The recommended approach to test the statistical significance of predictions in a cross-validation setting is to use a permutation test (Golland and Fischl 2003; Noirhomme et al. 2014). The outcome values are randomly permuted many times, and for each permutation, the cross-validation is performed using the permuted outcome values instead of original outcome values. A p-value is then calculated as a proportion of cross-validation results performed using the permuted data that is better than cross-validation results obtained using the original, non-permuted data.

A similar permutation testing procedure can also be used to obtain a null-distribution of an across cross-validation folds averaged confound adjusted test statistic e.g., ΔR^2^_p_ or ΔD^2^_p_ as described above. An important caveat is that the permutation procedure should only affect the relationship between input variables and the outcome, but not the relationship between the outcome and confounding variables (Fig 3) (Hoffling and Tibshirani 2008). The permutation needs to be performed on the rows of the input variables but not on the outcome labels and not on the confounding variables. If only the outcomes were shuffled, the results would be biased because the confounds will no longer be related to the outcomes, and thus this will not create a correct null distribution.

**Figure 3:**
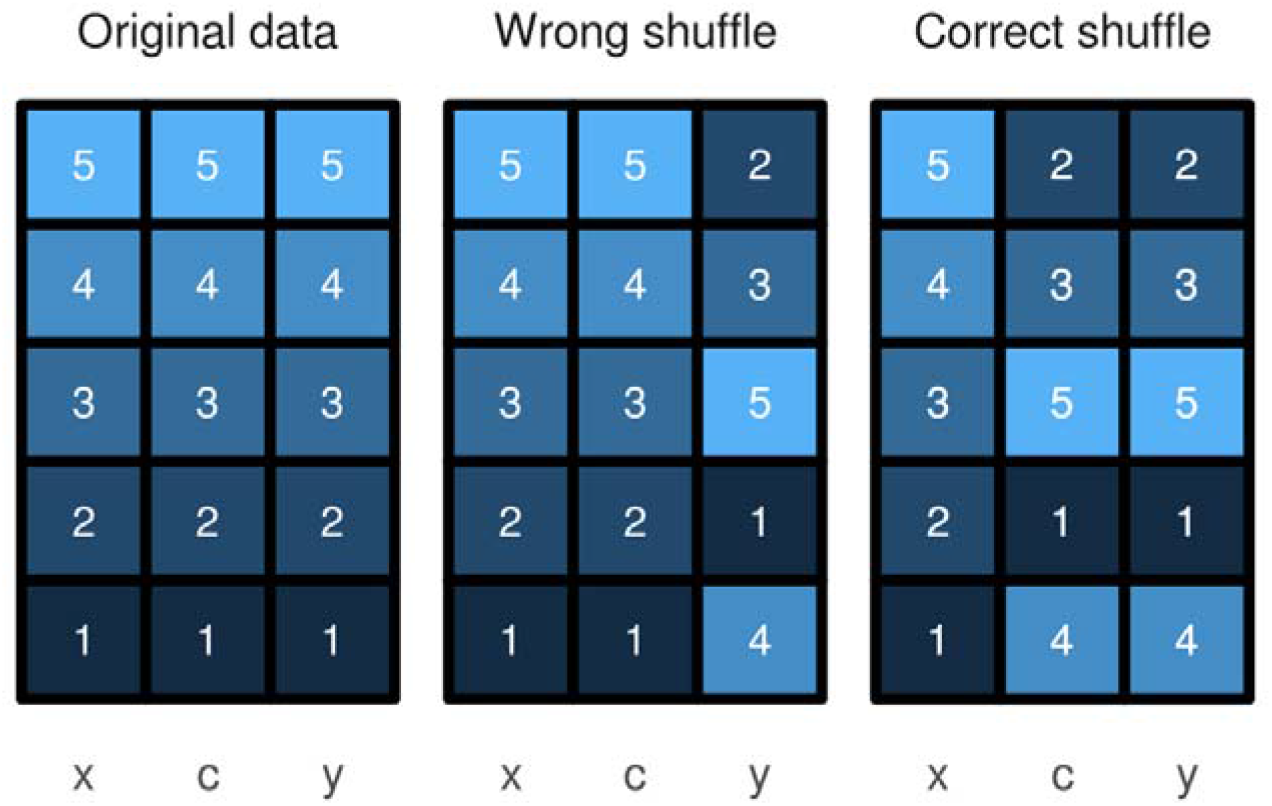
Valid and invalid permutation scheme in the presence of confounds. Given input variables x, confounds c, and outcome values y, the incorrect way is to shuffle only y, which would remove the relationship between x and y but also between c and y, leading to biased results. The correct way is to remove the relationship between x and y a but keep the relationship between c and y fixed.

## WHICH DATA TO USE TO CREATE THE ADJUSTMENT MODEL

The model used to perform confound adjustment can be estimated using all available data, however, in some cases, it has been recommended in the literature to use only a subset of the data to fit the confound adjustment model. Dukart and colleagues (2011) recommended performing confound adjustment only based on the data from healthy controls but omit data from cases, with the justification that otherwise we might be removing the effect of the disease as well, due to its interaction with the confound. However, as was pointed out by Linn et al. (2016), this procedure will not sufficiently remove the effects of confounds, and thus it will produce biased results as illustrated in Figure 4. This is because data from healthy controls are insufficient to estimate the effect of confounds in subjects with a disease.

**Figure 4:**
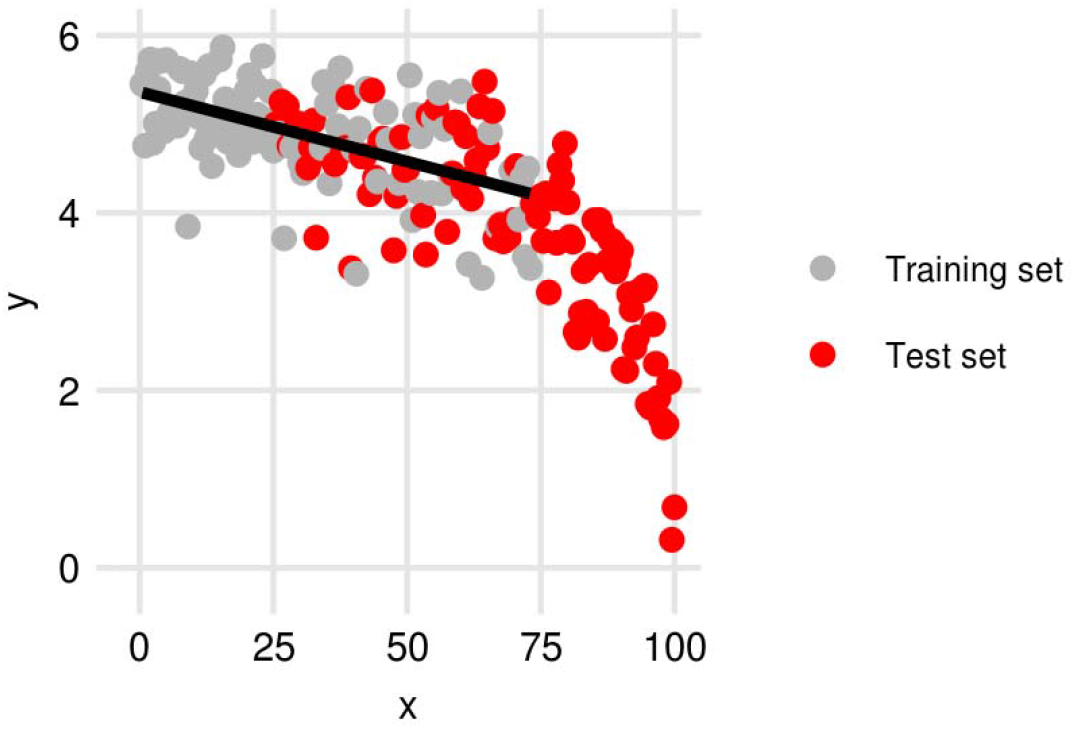
Adjustment of the test set based on training set data, or patients based on healthy controls data might be insufficient and thus should be avoided.

Snoek et al. (2019) recommend performing confound adjustment only based on the data from the training set but omit the test set to avoid a negative bias that can even lead to a significant below chance performance. If an effect of a variable on the outcome in the whole dataset is 0, then the effect learned in the training set will have an opposite sign in the test set, leading to negatively biased results.

To avoid this problem, Snoek et al. (2019) recommend that the confound adjustment of the test set data is performed using a model developed only on the data in the training sets, thus no artificial dependency between the training and test set data is created. However, in some situations, this is not possible because the levels of confounds we want to adjust for are not presented in the training data, for example, in the case of confounding effects of scan-sites where the test set consists of scan-sites that are not present in the training set. A second problem relates to a similar issue when adjusting only based on the data from healthy controls: if there is a difference in confound distribution between training and test set, then the test set will be insufficiently adjusted, and thus the results might be confounded. Since our proposed method is based on adjusting model predictions, it will not create an artificial dependency between the training and test set data, thus it will not lead to artificially lower than chance predictions.

## WHAT VARIABLES TO ADJUST FOR

Another important consideration is what variables we should adjust for. Since the goal is to evaluate machine learning predictive models, this is not necessarily a causal question, but mostly a practical one and thus entirely problem-dependent. We might want to ask two questions: 1) Is the machine learning model predicting a signal of interest or signals that are not of interest (e.g., scan-site, age, motion, BMI)? 2) Does the machine-learning model predict the actual outcome or perhaps something correlated to the outcome? For example, if we want to create a predictive model of future disease remission based on MRI data, we may want to ask whether our model based on MRI data predicts future remission or just current disease severity (which may be highly related to future remission). Therefore, we might want to control for current severity to estimate the added value of MRI data for prediction. However, maybe the goal of our study is to create an MRI based predictive model to replace a clinical assessment. In that case, the current severity of symptoms would not be considered a confound to control for, although we might still want to evaluate if the variance explained by severity is the same as the variance explained by the MRI based machine learning model.

## NONLINEAR AND NONPARAMETRIC ADJUSTMENT

The effect of confounds might not be strictly linear, thus it is important to be able to also accommodate nonlinear effects. If this is not considered, the residual effects of confounds can still be present in the data and can bias results (Figure 5). Since our proposed method is just an application of traditional linear or logistic regression, the same methods for estimating nonlinear effects apply. One way is to expand confounding variables using polynomial or basis spline expansions. Another possibility is to estimate the effects of confounding variables through non-parametrically generalized additive models and smoothing splines (Hastie and Tibshirani 1987; Wood 2017), according to the following equation:

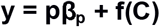

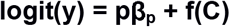

**Figure 5:**
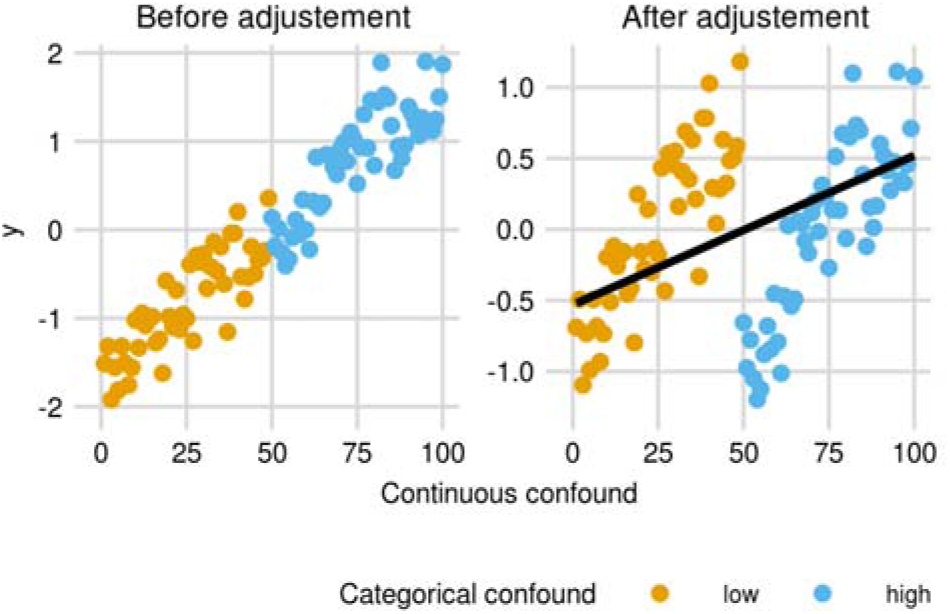
Categorizing continuous confound variable before adjustment might lead to insufficiently adjusted data, with the residual confounding signal still present in the data.

Where **f(C)** is a parametric or non-parametric smoothing function.

A somewhat common, but invalid approach to account for nonlinear effects of confounds is categorizing confounding variables. For example, instead of correcting for BMI, the correction is performed for categories of low, medium, and high BMI. Such a categorization is unsatisfactory because it keeps residual confounding within-category variance in the data, which can lead to both false positive and false negative results (Austin and Brunner 2004). False-positive results because there can still be residual confounding information presented in the input data, and false negative because the variance in the data due to confounding variables will lower the statistical power of a test. Thus, categorizing continuous confounding variables should not be performed. Instead, other parametric or nonparametric approaches for the modeling of nonlinear effects should be used.

## EXAMPLE ANALYSIS

To illustrate the usage of the proposed approach, we performed an example analysis in a similar way that it can be performed in practice. We aimed to predict a fluid intelligence score (FI) based on volumetric data of brain regions of interest. We performed a separate control for two confounding variables, 1) brain size, to evaluate if a machine learning model learned any useful patterns from the data, above and beyond what can be explained by brain size. 2) Age, when a continuous full time education was completed, to evaluate the added benefit of a brain-based machine learning model to a variable that is much easier to obtain and test if the ML model learned to predict anything else than function of education length.

### Dataset

We used the 2018 release of the UK Biobank dataset, specifically region of interest volumetric data as obtained from the structural MRI images using FSL FAST. The details about the dataset, preprocessing, and volume extraction procedure can be found elsewhere (Sudlow et al. 2015; Fawns-Ritchie and Deary 2020; Miller et al. 2016; Alfaro-Almagro et al. 2018). Prior to analysis, we used the funpack utility v2.0.1 (McCarthy 2019) to compute imaging derived phenotypes and non-imaging data. From the dataset of size n=21,407 we excluded 2,149 subjects due to missing FI data. Next, we split the data into halves while preserving the FI distributions between a training and test set. We also excluded 4,461subjects from the test set due to missing education data, leaving the size of the training set at 9,630 and the test set 5,167.

### Method

We fitted a ridge regression model implemented in the glmnet package (Friedman et al. 2010) in the training set to predict FI using the regional brain volumetric data. Next, we used this model to obtain predicted FI scores for subjects in the test set. We evaluated the predicted test set FI scores by including them in two multivariable linear regression models with the brain size variable (model 1) or with an age of completed full-time education (model 2) as covariates. We tested if the predicted FI scores are statistically significant in these models and estimated their partial R2 given covariates. To take into account nonlinear effects of education, we used cubic spline expansion with 5 knots. This procedure allowed us to estimate the proportion of the FI, explained by confounding variables, and a proportion of FI variance explained by predictions alone, thus effectively controlling the effects of confounding variables. Note that the machine learning model was built in the training set, but statistical tests were performed in the test set.

### Results

Machine learning predictions, e.i., the predicted FI score based on the volumetric data, explained a small proportion of FI variance (R^2^=0.035, p < 0.001), which could partly be explained by brain size (R^2^_shared_ = 0.014; ΔR^2^_MLpredicitions_ =0.021, p < 0.001). This indicates that ML can learn patterns related to FI above and beyond brain size, even though the added benefit is small. Brain size was not able to explain additional variance not already explained by ML predictions (ΔR^2^_brainsize_ = 0, p=0.290), which was expected since the effect of brain size can be implicitly learned from the individual region of interest volume measures. The R^2^ due to ML predictions (R^2^_ml_predicitions_ = 0.039, p < 0.001) was smaller than R^2^ due to education length (R^2^_education_=0.079, p < 0.001). However, the ML predictions explained a proportion of variance that could not be explained by the effect of education. Therefore, the partial effect of ML predictions was still highly statistically significant.

## EXPERIMENTS

For the empirical validation of the proposed method, we simulated confounded datasets for regression and classification and also used a real dataset with the artificially created confounded outcome variable. The dataset for regression was simulated similarly to the dataset in Fig 2B. The confounding variable was uniformly distributed between 0 and 100. Input variable x was simulated as a function of the confound with heteroscedastic noise term according to:

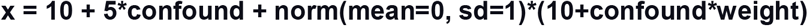

where the weight was set to 0.1, 0.2, and 0.3 representing low, medium, and high effect of confounding. The outcome variable y was simulated as a function of the confounding variable according to:

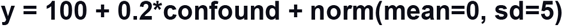

therefore, there was no relationship between x and y that cannot be explained by the effect of the confounding variable.

The dataset for classification was simulated similarly to Fig 1C. First, a categorical confounding variable with two categories was created. Next, two input variables x1 and x2 were sampled from a bivariate normal distribution with mean = 0, 0 and covariance matrix = 1 0.9 0.9 1 or 1 −0.9 −0.9 1, depending on the category of the confounding variable. Outcome variable y was created as a function of the confounding variable, such as the probability of y=1 was 0.6, 0.7, 0.8 representing low, medium, and high confounding, if the confounding variable was 0.

We use support vector regression with a radial basis function (RBF) kernel with C=1 for the regression problem (Smola and Schölkopf 2004) and Gaussian process classification (Rasmussen and Williams 2005) with an automatic sigma estimation for the classification problem, both implemented in kernlab (Karatzoglou et al. 2004). For each simulation, we created a dataset of size 200. We assessed the statistical significance of machine learning predictions in two ways: 1) in the test set, by splitting the dataset into halves used as a training set and a validation set where the statistical significance was assessed using a parametric test. 2) in a 5-fold cross-validation, where the statistical significance was assessed using a permutation test. Our goal was to adjust for the confounding variables. We performed two types of confound adjustments. 1) adjustment of input variables according to (Snoek et al. 2019), where the training set data and test set data were first adjusted for confounds by regressing out the effect of confounding variables estimated using the training dataset. 2) additional adjustment of the ML predictions according to the proposed method using linear regression for the regression problem and logistic regression for the classification problem. The statistical significance was assessed either parametrically on the holdout set, or nonparametrically using a permutation test for cross-validation. Since the datasets were created, such as there is no relationship between input variables and the outcome that is not explained by the effect of confounding variables, the valid confound adjustment method should produce statistically significant results with p < 0.05 around 5% of the time.

We also used a real neuroimaging dataset consisting of region of interest (ROI) thickness data from the ABIDE dataset (Di Martino et al. 2014) obtained from the preprocessed connectome project (Craddock et al. 2013). The dataset consisted of 97 regions of interest (ROI), including cortical ROI obtained using the ANTs pipeline according to Desikan-Killiany protocol (Klein and Tourville 2012) and FreeSurfer white matter and non-cortical ROI (Fischl 2012)(See https://mindboggle.readthedocs.io/en/latest/labels.html for a complete list of ROIs). We used data from 530 healthy control subjects, with age at scan spanning between 6 to 56 years (median = 14.66, IQR=8.2).

For each simulation, we randomly created a categorical outcome variable y confounded by age, where the probability of y=1 was based on subjects age according to:

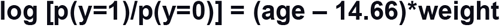

where weight was set to 3, 4, and 5 representing low, medium, and high confounding, since the outcome variable was created solely as a function of age, there should be no signal in the data after adjustment for age.

## RESULTS

Since the simulated datasets were created so that there is no signal in the data that cannot be explained by the confounding variables, all statistically significant results are false positive. Therefore, the successful de-confounding method should have a false positive rate at p < 0.05 around 5%. In simulated regression and classification datasets, the input adjustment method did not sufficiently control for confounds, and significantly above chance performance was obtained. The percentage of false-positive results could be made arbitrarily high (even as high as 100%), depending on the amount of confounding. In the real dataset, a false positive rate as high as 72% was observed when predicting the outcome that was created as a function of age after correcting for age. In all tested scenarios, our proposed confound adjustment method of the machine learning predictions achieved a false positive rate of around 5%. This was the case when the testing was performed on the independent hold-out set using a parametric test or in cross-validation using a permutation test (Figure 6).

**Figure 6:**
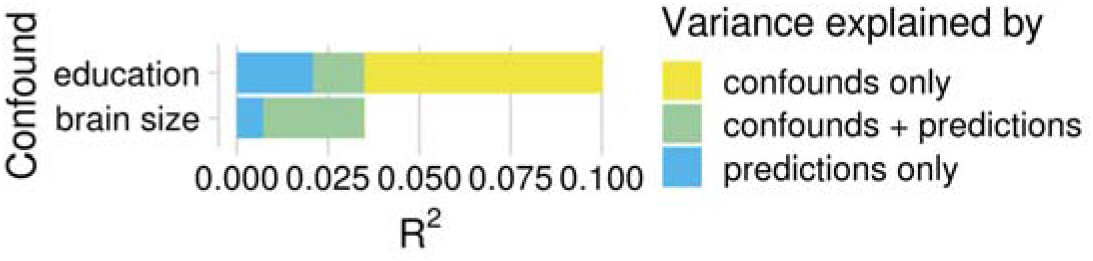
Results of machine learning prediction of fluid intelligence based on brain imaging data, taking into account the confounding effects of education length or brain size. Machine learning predictions were able to predict a proportion of variance not already explained by the impact of confounding variables, therefore the results were not fully driven by confounds.

**Figure 6:**
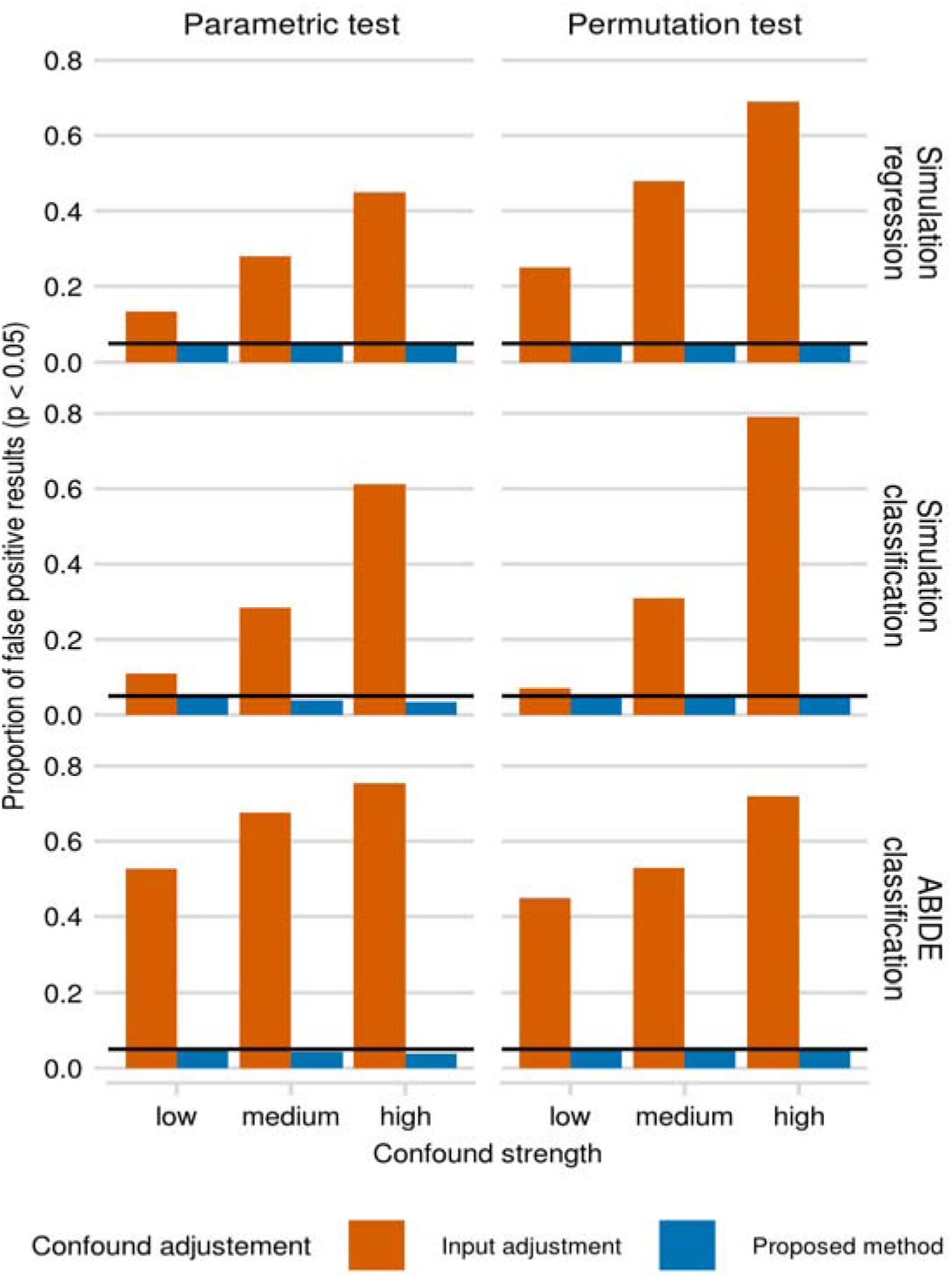
Results of traditional input adjustment method and proposed output adjustment method. The datasets were created so that there is no signal in the data that cannot be explained by the effect of confounding variables, therefore after successful confound adjustment, the proportion of statistically significant results (p < 0.05) should be around 5% (black line). We simulated confounded data for regression and classification problems. Also, we used a real neuroimaging dataset (ABIDE), where the confounding variable was age, and we created an artificial categorical outcome as a function of age. The amount of confounding effect was varied. The statistical significance was assessed using a parametric test in the hold-out set, or a permutation test in 5-fold cross-validation.

## CODE AVAILABILITY

The analysis code is available at https://github.com/dinga92/confounds_paper

## DISCUSSION

We demonstrated that controlling for confounds at the level of model predictions correctly adjusts for the effect of confounding variables under low, medium, and high confounding, even in situations where the traditional input variable adjustment failed. Moreover, this approach has a simple and intuitive interpretation: what proportion of variance in the outcome can be explained by model predictions that cannot be explained by confounding variables? Or in other words, does predicting the outcome using confounding variables improve when adding model predictions? The proposed approach is based on testing if machine learning predictions are statistically significant in a linear or logistic regression model that includes confounding variables as covariates. The proposed approach evaluates if the model predictions have any information about the outcome that is not already present in the confounding variables.

Traditional input variable adjustment failed to sufficiently control for confounds in simulated and real datasets. This is because input variable adjustment cannot remove all confounding effects that can be learned by machine learning methods, as we show illustrative examples and in the simulated data. This includes cross-validated input adjustment as proposed by (Snoek et al. 2019) and adjustment using a location and scale adjustment model as used in ComBat (Fortin et al. 2017). Therefore, it is possible that some of the previously published machine learning results are (partly) driven by insufficiently adjusted confounding instead of the signal of interest. Machine learning methods vulnerable to this problem include all nonlinear machine learning methods and linear machine learning methods that are fitted optimizing a different function than a regression used for input adjustment, such as support vector machines. Support vector machines optimize a hinge loss, which is more robust to extreme values than a squared loss used for input adjustment. Therefore, the presence of outliers in the data will lead to improper input adjustment that can be exploited by SVM. Studies using penalized linear or logistic regression (i.e., lasso, ridge, elastic-net) and classical linear Gaussian process modesl should not be affected by these confounds (assuming they are sufficiently adjusted) since these models are not more robust to outliers than OLS regression.

The proposed approach is closely related to the “pre-validation” method used in microarray studies to test if a model based on micro-array data adds value to clinical predictors (Tibshirani and Efron 2002; Hoffling and Tibshirani 2008). Here we argued that this approach can also be used to control for confounds of machine learning predictions in general and suggest using D^2^ and R^2^ and their decompositions to interpret the results. Multiple alternative approaches for controlling confounding effects exist and can be used in a machine learning setting. A few indirect methods have been proposed. For example, Görgen et al. (2017) proposed to test whether a machine learning model can accurately predict confounding variables, and Kohoutova et al. (2020) proposed to test whether confounding variables can predict the outcome, thus diagnosing potentially spurious results. Our proposed method's benefit is that it provides a direct formal test of the association between machine learning predictions and the outcome controlled for confounds.

A related approach to the proposed method, sometimes used in neuroimaging (for example in ABCD Neurocognitive Prediction Challenge 2019 (Pohl et al. 2019)), is to residualize target variables, i.e., regress out confounds from the target variables and apply machine learning to predict these residuals. This approach is similar to testing for partial correlations, as we proposed, with a few critical distinctions. First, the partial correlations are equivalent to regressing out confounding variables from both target variable and ML predictions. Thus, this approach accounts for more variance in the data, making it more sensitive and increasing statistical power. Second, regressing out confounds from the target variables cannot be applied to classification problems. Third, the interpretation is limited, since it only allows for testing the partial effects of ML predictions, while our proposed method allows for the estimation of variance that can be explained by both confounds and machine learning predictions, confounds only, and machine learning predictions only. Fourth, the target residualization is often applied to the whole dataset, which leads to data leakage between the training set and the test set and thus to biased results (Snoek et al. 2019). This data leakage can be avoided by estimating model parameters using only training set data, however, this might also lead to biased results due to insufficient confound adjustment in the test. In contrast, the proposed approach is applied only in the test set, which avoids the data leakage and ensures that the effect of confounds is sufficiently estimated.

Controlling for confounds can also be done using a permutation test where the permutations are performed within the confound groups (Winkler et al. 2015). For example, if we wish to control for effects of scan sites, labels would be shuffled within each scan site separately. Thus if a model‘s performance is driven by the scan site effects, this will be reflected in the permutation-based null-distribution. Although this method can be used even with a complex dependence structure (Winkler et al. 2015), it is limited to situations where the dependence structure is clear, and includes only a few categorical confounding variables, since the number of confounding variables limits the number of possible permutations.

Another possibility is to use various resampling or reweighting methods to create a dataset where the confounding variable is not related to the outcome (Pourhoseingholi et al. 2012; Rao et al. 2017; Chyzhyk et al. 2018). Since only a subset of available subjects is used, this leads to data loss and highly variable estimates. Another problem of this approach is that the distribution of variables in the test set no longer matches the distribution of the original dataset or the population. For example, when controlling for a sex effect in the machine learning prediction of Autism diagnosis, resampling methods would be interpreted as the performance of the machine learning model in a population where sex is not related to the autism diagnosis. Since more males are diagnosed with autism than females, resampling methods will put a higher weight on predictions for females, which might lead to the selection of a model that makes worse predictions in males even though the majority of autism diagnoses are in males.

A related problem of resampling methods is that they cannot be used to evaluate the additional benefit of the machine learning model predictions over what can be already predicted by confounds since by definition confounding variables have no predictive power in the newly created population. It might be tempting to say that the model's added value equals the performance of the model in this newly created population. This is, however, not the case. As shown by Pepe et al. (2012) and Janes and Pepe (2008), this can severely underestimate and also overestimate the added value and even change ranks of competing models. Thus, it can lead to selecting the worse model for prediction, missing potentially important biomarker, or selecting an apparently strong biomarker that, in reality, does not add much to what can be already predicted using confounds.

## CONCLUSION

We showed that confound adjustment of input variables can fail to adequately control for confounding effects when machine learning methods are used. For this reason, we propose that confound adjustment of input variables should be avoided, and the already published machine learning studies employing this method should be interpreted with care. We presented a simple approach of controlling for confounds at the level of machine learning predictions themselves. This approach produced more valid results even under heavy and complicated confounding.

